# Interpretable Drug Response Prediction through Molecule Structure-aware and Knowledge-Guided Visible Neural Network

**DOI:** 10.1101/2024.02.07.579280

**Authors:** Jiancong Xie, Zhe Zhang, Youyou Li, Jiahua Rao, Yuedong Yang

**Affiliations:** School of Computer Science and Engineering, Sun Yat-sen University, Guangzhou, 510000, Guangdong, China; Department of Chemistry, University of Chicago, Chicago, 60637, Illinois, USA; Key Laboratory of Machine Intelligence and Advanced Computing of MOE, Sun Yat-sen University, Guangzhou, 510000, Guangdong, China

**Keywords:** Drug response prediction, signaling pathway, transcriptome, message passing network, visible neural network

## Abstract

Precise prediction of anti-cancer drug responses has become a crucial obstruction in anti-cancer drug design and clinical applications. In recent years, various deep learning methods have been applied to drug response prediction and become more accurate. However, they are still criticized as being non-transparent. To offer reliable drug response prediction in real-world applications, there is still a pressing demand to develop a model with high predictive performance as well as interpretability. In this study, we propose DrugVNN, an end-to-end interpretable drug response prediction framework, which extracts gene features of cell lines through a knowledge-guided visible neural network (VNN), and learns drug representation through a node-edge communicative message passing network (CMPNN). Additionally, between these two networks, a novel drug-aware gene attention gate is designed to direct the drug representation to VNN to simulate the effects of drugs. By evaluating on the GDSC dataset, DrugVNN achieved state-of-the-art performance. Moreover, DrugVNN can identify active genes and relevant signaling pathways for specific drug-cell line pairs with supporting evidence in the literature, implying the interpretability of our model.

## 1. Introduction

Cancer constitutes an immense challenge to human health, necessitating a significant demand for effective anticancer medications[1]. Although each year thousands of new medicines are developed for the treatment of cancer, fewer than 4% are ultimately approved by the US Food and Drug Administration[2, 3]. One of the main reasons for this low approval rate is an incomplete understanding of the mechanisms of new candidates in cells. With this respect, an accurate and interpretable drug response prediction (DRP) model is vital in the drug discovery process to facilitate novel and practical cancer treatment development.

Recently, the accumulation of cancer multi-omics data provides an opportunity to comprehend tumors’ omics characteristics and their effects on drug responses[4]. Driven by the advances in high-throughput technologies, several large-scale cancer genetic projects have been launched and systematized into public repositories, including Genomics of Drug Sensitivity in Cancer (GDSC)[5] and Cancer Cell Line Encyclopedia (CCLE)[6]. These public databases provide a wealth of genetic profiles of cancer cell lines and drug response information leading to the development of DRP techniques[7].

Early studies in DRP utilized machine-learning techniques like random forest [8], support vector machine[9], and matrix factorization[10]. Although these early attempts have made remarkable progress, their simplistic model structures struggle to capture the high-dimensional representation of drug and omics data, resulting in poor generalizability. In recent years, with the capabilities of learning complex nonlinear functions, various deep learning techniques for DRP have garnered substantial attention[11, 12, 13]. CDRscan[11], for instance, firstly introduced a 2D-convolution model by combining molecule fingerprints as drug features and genomic profiles as cell features. MOLI[12] further improved the prediction accuracy using a type-specific sub-network to encode the multi-omics data. However, these deep learning methods for DRP serve as “black boxes”, lacking interpretability in terms of biological mechanisms[14], while a rational explanation of a new drug’s biological functions is necessary for its further clinical application.

To improve interpretability, recent studies sought to encode biological knowledge directly into the architecture of the neural network by mapping the neurons into a specific biomarker[15, 16]. These models were named visible neural networks (VNNs)[17] as opposed to the traditional fully connected deep neural networks (DNNs). As a representative example, PathDNN[18] reshaped the canonical DNN structure by incorporating a layer of pathway nodes and their connections to input gene nodes. Similarly, DrugCell[17] embedded the Gene Ontology[19] and Clique-eXtracted[20] Ontology into the architecture of the neural networks and encoded gene deletion genotypes via VNN for predicting drug responses. Alhough these methods show excellent simulation and interpretability, the predictive performance is limited due to their incapability of learning drug geometry information.

In fact, molecules can be represented as graphs where atoms are nodes and bonds between atoms are edges. The graph representation for molecules has been shown to outperform designer representations like fingerprints in many tasks[21]. Recently, graph-based approaches have been introduced in DRP[22, 23, 24]. For example, DeepCDR[23] proposed a uniform graph convolutional network to capture the intrinsic chemical structures of drugs, and DeepTTA[24] adopted a transformer architecture to mine the sub-structure features of compounds. Taking advantage of the structural information of drugs, these methods outperform PathDNN and DrugCell by a large margin. Overall, high accuracy and interpretability are both necessary for drug response prediction models, but current methods have drawbacks either in the interpretability of the mechanism of drug action or limited performance in modeling drug response. Moreover, the abovementioned methods are basically two-branch networks where drug and cell line encoders extract their own features separately. When considering the interaction of drugs with their targets leading to cells’ physiological and biochemical changes, they fail to learn the intrinsic effect of drugs on cells.

To address the limitations above, we proposed a novel DRP framework, DrugVNN, to combine the powerfully expressive graph convolution network with an interpretable knowledge-guided visible neural network. As inspired by the previous work[25, 26], we build a node-edge communicative message passing attention network, with local bond-aware attention mechanism during message passing process and global attention pooling during readout process. In parallel, we construct a cancer-related pathway VNN, where the neuron units are sparsely connected based on prior pathway knowledge. Between these two networks, a drug-aware gene attention gate is designed so that each gene is assigned different weights using QKV Attention[27] based on the learned drug embedding to select useful, relevant gene transcriptome information. The learned drug and cell line embedding are finally concatenated to predict the IC_50_. By comparison with the existing DRP methods, DrugVNN achieved state-of-the-art performance and successfully identifies the most relevant pathways driving the prediction, providing evidence for the underlying mechanisms of the drug responses.

## 2. Materials and methods

### 2.1. Data preparation

#### 2.1.1. Drug response data

The drug response data was collected from the GDSC2 database[5], which provides log normalized IC_50_ values across different drugs and cell lines. By excluding the drugs without Pub-Chem ID and cell lines in which any type of omics data was missing, a total of 123,760 IC_50_ pairs spanning 204 drugs and 684 cell lines were finally selected. All the cell lines belong to 9 cancer types. We further downloaded the simplified molecular input line entry system (SMILES) of each drug from PubChem database[28]. Among all the 204 × 684 = 139, 536 drug-cell line interactions pairs, approximately 11.3% (15,776) of the IC_50_ values were missing.

#### 2.1.2. Genomic data

The multi-omics profiles of cancer cell lines were collected from the CCLE database[6], where three types of gene-level omics data were considered: gene expression, somatic mutation, and copy number variation. The expression data were RNA sequencing (RNA-seq) log2-transformed TPM (transcripts per million) values. The copy number variation was log2-transformed with a pseudo-count of 1. The somatic mutation data were represented as a binary vector where “1” indicated that the gene was mutated. To exclude the low-expressed genes, we only considered the genes with expression values greater than 1 in at least 1% of the cell lines. This yielded 8542 genes and the input of each cell line is represented by a 3 × 8542-dimensional feature vector.

#### 2.1.3. Pathways data

To design the VNN architecture that simulates the cells’ biological process, we collected the pathway gene set and hierarchy relationship files from Reactome (https://reactome.org/)[29] and selected the pathways with at least five genes as feature genes. For the pathway relation *P*_1_ → *P*_2_, *P*_1_ was denoted as the child and *P*_2_ as the parent. A root node was assigned as the parent node for the pathways without parents to aggregate high-level pathway information. Consequently, we obtained 1774 pathways with 13 layers, where pathways directly associated with the input genes were at the lowest level (layer one) and the root node was at the highest. A profile of the basic statistics of the pathway data is shown in Supplementary Table S1.

### 2.2. The architecture of DrugVNN

In this study, we developed a regression model to predict the half maximal inhibitory concentration IC_50_, a representative indicator of drug sensitivity. Fig. 1 shows the overall architecture of DrugVNN with two branches: drug feature encoder modeling molecule geometry information and cell line feature encoder modeling the hierarchical organization of molecular subsystems in the cell. Two branches interact through a drug-aware gene attention gating mechanism.

**Fig. 1.**
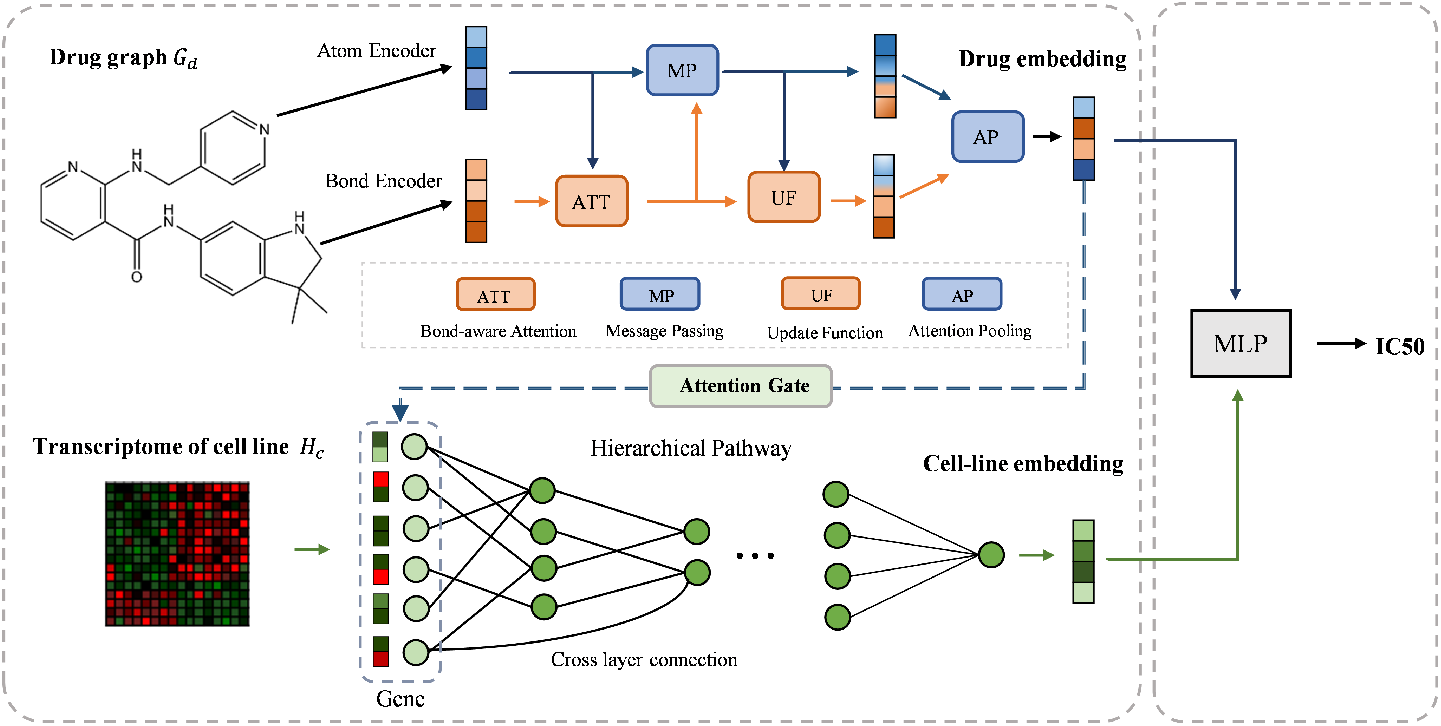
Overview of the DrugVNN framework. DrugVNN takes a pair of drug and cancer cell line profiles as inputs. In the drug representation learning section, the initial features of atoms and edges are fed to the node-edge communicative message passing module and the final embeddings are weighted averaged among different atoms based on attention pooling mechanism. In cell line representation learning section, the learned drug embeddings first interact with the input gene profiles to select useful gene information, which is then fed into a hierarchical pathways-guided VNN to extract the final embedding of the cell line.

#### 2.2.1. Drug feature mining by CMPNN

Graph representation learning has been extensively applied to molecule feature mining, where molecule atoms are represented as nodes with bonds between atoms represented as edges. Message Passing Neural Network (MPNN)[30] is one of the most widely used graph representation learning methods, which learns node representations by aggregating neighbor information during the message-passing stage. In the readout stage, MPNN utilizes a permutation-invariant readout function to summarize all node representations into graph-level representations.

To enhance the ability of MPNNs that only consider node-node message passing, node-edge communicative MPNNs (CMPNN)[25] were proposed and achieved state-of-the-art performance in molecule representation learning. Following the key idea of CMPNN, we adopted the node-edge interaction module and further designed a local attention mechanism in the message passing process and global attention in the pooling process to improve the learning ability of the model.

Formally, given a directed molecular graph *G* = (*V, E*), the initial atom features *H*_*v*_ and edge features *H*_*e*_, We first mapped the node and edge representations to the same dimensionality *f*:

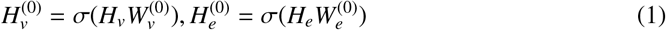

where 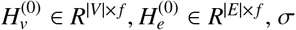 denotes the activation function, 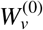 and 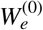 are learnable parametric matirx. Then, L-layer propagations were applied.

##### Bond-aware attention

In layer *l*, before message passing, the attention mechanism was employed to optimize node information aggregation. The bond embedding and its connected atom embedding were used to calculate the weight of each bond, which provided dynamic weighting capability and enabled the model to eliminate redundant information in message aggregation.

The attention scores were calculated as follows:

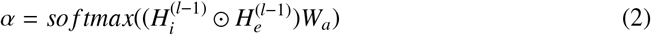

where *H*_*i*_ ∈ *R*^|*E*|× *f*^ represents the source nodes embedding of the edges, *W*_*a*_ denotes the learnable parametric matrix, and the operator ⊙ denotes the elements-wise product. Then we obtained the attentive bond embedding:

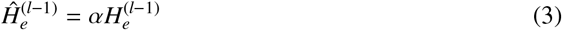

##### Node message passing

In the message passing module, the node embeddings were updated by aggregating the attentive embeddings of their incoming edges:

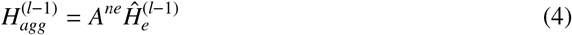

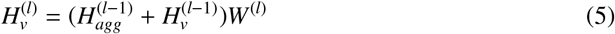

where *A*^*ne*^ is the node-to-edge adjacency matrix that connects each node to its incoming edges. To further improve the learning ability, we used K different graph convolution kernels at each layer to extract local features of different sizes separately:

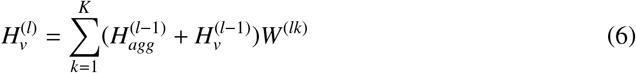

where *W*^(*lk*)^ denotes the learnable parametric matirx of *k*^*th*^ kernel in *l*_*th*_ layer.

##### Bond embedding updating

Most current MPNNs only updated node embedding and ignored the information carried by bonds. Herein we updated bond embedding in every iteration using the learned atom embedding:

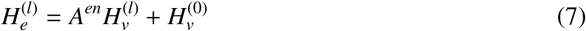

where *A*^*en*^ is the edge-to-node adjacency matrix that connects each edge to the source nodes.

##### Attention pooling

After *L* layers, instead of using the global mean pooling that might either dilute or lose the important features, we employed self-attention pooling to automatically focus on important atoms. Specifically, we obtained the self-attention matrix *A* ∈ *R*^|*V*|×*n*^ with *n* attention heads:

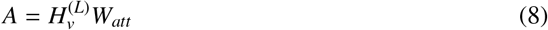

where *W*_*att*_ is learnable parameters. Then the graph-level representation was calculated as follows:

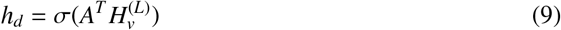

By continuously updating and fusing the atomic and bond features, we obtained the final drug embedding *h*_*d*_ for further incorporation in cell line representation learning and drug response prediction.

#### 2.2.2. Cell line feature mining by Pathways-guided VNN

Drug molecules interact with their targets and subsequently inhibit or activate relevant pathways that control the cellular processes or influence tumor microenvironments such as cell proliferation, angiogenesis, and immune response. To simulate these biological processes, we integrated gene–pathway and pathway–pathway relationships data and built the hierarchical pathway-guide visible neural networks (VNN).

Specifically, we mapped each neuron of the neural network into different pathways. Neurons are connected only when an association exists between the corresponding pathways. The input of VNN consists of the genomics profile of the gene, including expression, copy number and mutation as described in the data preparation section. The first hidden layer of the lowest-level pathways selectively connects the input genes to the neurons in the same pathways. Then, it sends the integrated information to the neurons of corresponding parent pathways until reaching the root. At the end of the VNN, the output of the root pathway is extracted as the final embedding of the cell line.

Mathematically, we denoted the set of genes as *G* = {*g*_0_, *g*_1_, …, *g*_*n*_}, and the genomics profile of gene as *h*_*g*_, where *n* is the number of genes. The input of each pathway (neuron) *p* consists of two parts:

- The output of its child pathways, which was concatenated into a single vector, denoted as *O*_*p*_.
- The genomics profile *h*_*g*_ of its connected genes, which was concatenated into a single vector, denoted as *H*_*p*_.

Notably, the pathway units in the first layer of VNN contain only genomic features as input. Then we obtained the output of pathway *p* as follows:

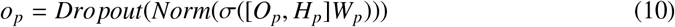

where *W*_*p*_ is the learnable parameters of pathway *p*, σ, *Norm* and *Dropout* is the activation, normalization, and dropout function. In the last layer of VNN, the output of the root pathway, integrating the information of all other low-level pathways, is obtained as the final embedding *h*_*c*_ of the cell line.

#### 2.2.3. Drug-aware gene attention gates

Drugs interact with specific targets to play their functions in cellular activities. To simulate the mechanism of drug action, we proposed a drug-aware gene attention gates mechanism to select useful genomic information based on the learned embedding of the drugs. In this way, we expected our VNN to realize which target is more active for a specific drug. Here, we adopted the QKV attention[27] mechanism between the learned drug embedding and gene genomic profile, where the drug is treated as a query, while the genes in cancer cells are attended to as keys and values:

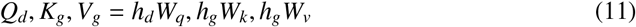

where *W*_*q*_, *W*_*k*_, *W*_*v*_ are learnable matrices. Then, QKV Attention is applied to {*Q*_*d*_, *K*_*g*_, *V*_*g*_} to extract beneficial information and update the representation of genes:

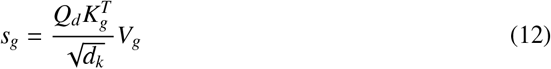

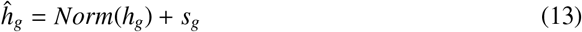

The updated gene representations are then input to the VNN, guiding the network to better simulate the biological process of the drug on the cell line.

For interpreting the learned VNN model, we aim to extract a high-attribution sub-network consisting of contributing pathways that are highly relevant for drug response prediction. To achieve this goal, we adopted SHAP (SHapley Additive exPlanations)[31], a game theoretic approach for interpreting the predictions of a DNN by generating prediction values (SHAP values) for each feature. A higher SHAP value stands higher contribution of a feature to the prediction result.

Due to the hierarchical structure of VNN, we extracted the high-attribution sub-network in a top-down manner. Given a drug to interpret, we first computed the SHAP values *s*_*p*_ of each node *p* in the hidden layer in VNN and set a threshold τ. Then we identified the important nodes set *V*_*l*_ in layer *l* by following the rules: (1) *s*_*v*_ > τ for each node *v* ∈ *V*_*l*_, and (2) more than one parent pathways u of v that *u* ∈ *V*_*l*+1_. In the beginning, we initialed *V*_*L*_ = {*root*} and iterated the above selecting process up to the lowest-level pathways. The final high-attribution sub-network consists of the identified important nodes and edges between these nodes.

## 3. Results

### 3.1. Experimental Design

To verify the effectiveness of DrugVNN, we compared our model with other six state-of-the-art DRP methods, including CDRscan[11], tCNNs[13], PathDNN[18], DrugCell[17], DeepCDR[23] and DeepTTA[24]. We evaluated our method under two experimental settings. (i) For re-discovering known drug-cell line responses: the whole dataset was randomly split into training/validation/test sets with a ratio of 8:1:1; (ii) For blind test (leave-drug/cell-line-out): the whole dataset is split on the drug or cell line level to guarantee that the test set only includes unseen drugs/cell lines in the training stage with the same split ratio of 8:1:1. We also designed a classification task by predicting binary drug sensitivity status. For model optimization, we utilized the Adam optimizer with weight decay of 10^−5^and the learning rate of 1 × 10^−4^. The proposed method was implemented with Pytorch 1.13.0 on an Nvidia GeForce RTX 3090 GPU.

Similar to the previous works, we evaluated the regression prediction of our model by root mean square error (RMSE), mean absolute error (MAE) and R-squared (R^2^) as the following definitions:

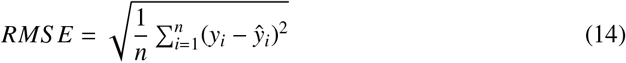

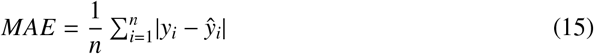

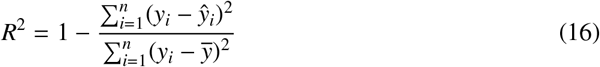

where *n* represents the number of samples, 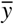 is the mean of the ground-truth data. For the classification task, AUROC and AUPR are adopted for evaluation.

### 3.2. Predicting drug-cell line responses

We first compared our proposed model with the state-of-the-art baselines by rediscovering known drug-cell line responses. All possible drug-cell line pairs were randomly divided into training, validation, and test sets in an 8:1:1 ratio. As shown in Table 1, our proposed method DrugVNN achieved the best performance on all three metrics with RMSE of 0.935, MAE of 0.685 and R^2^ of 0.885. We noticed that two VNN-based baselines (PathDNN and DrugCell) surpassed CDRscan and tCNNs. This is expected as VNN incorporates prior domain knowledge to simulate the biological processes inside cells and generates more reasonable features. We also noticed that the GNN-based methods (DeepCDR and DeepTTA) achieved superior performance over other baselines, which is consistent with the observation that GNNs tend to outperform traditional methods on many molecule-related tasks[32]. In DrugVNN, both GNN and VNN are employed to achieve outstanding predictive performance.

**Table 1.** Comparison of the model architectures and performances.

| Method | Feature Encoder |  | RMSE | MAE | R<sup>2</sup> |
| --- | --- | --- | --- | --- | --- |
|  | Drug | Cell line |  |  |  |
| CDRscan | CNN | CNN | 1.349±0.012 | 0.982±0.014 | 0.753±0.001 |
| tCNNs | CNN | CNN | 1.275±0.013 | 0.952±0.014 | 0.788±0.002 |
| PathDNN | - | VNN | 1.258±0.016 | 0.938±0.011 | 0.791±0.003 |
| DrugCell | FCN | VNN | 1.164±0.021 | 0.862±0.015 | 0.819±0.003 |
| DeepCDR | GNN | FCN | 1.082±0.020 | 0.791±0.017 | 0.842±0.002 |
| DeepTTA | GNN | FCN | 1.025±0.008 | 0.752±0.007 | 0.865±0.001 |
| DrugVNN | GNN | VNN | **0.935±0.006** | **0.685±0.006** | **0.885±0.001** |

To further examine the generalization ability of our method, we compared the proposed model with the best baseline DeepTTA by blind test settings, where drugs or cell lines in the test set were unseen during the training process. As shown in Fig. 2A and 2B, although the performance of our method and DeepTTA both decreased due to the out-of-distribution issue, DrugVNN still outperformed DeepTTA in both blind test scenarios. Specifically, in the leave-drug-out scenario, DrugVNN achieved slightly higher average Pearson’s correlation coefficient (0.667) than DeepTTA (0.650). 13 of 20 (65%) unseen drugs were predicted more accurately by DrugVNN. In the leave-cell-out scenario, DrugVNN outperformed DeepTTA by a large margin. Among the total 68 cell lines, 64 (94%) were more accurately predicted by DrugVNN. We illustrated a prediction case of cell line ZR7530 in Fig. 2C and 2D, where DrugVNN achieved a significantly higher Pearson’s correlation coefficient (0.833) than DeepTTA (0.636). The improvement can be attributed to obtaining better cell line representation through the knowledge-guided VNN. To demonstrate the effect of VNN, we visualized the t-SNE[33] 2D representation of the cell lines from the top-5 frequent tissue categories. As shown in Fig. 3, DrugVNN can better distinguish cell lines from different tissues than DeepTTA.

**Fig. 2.**
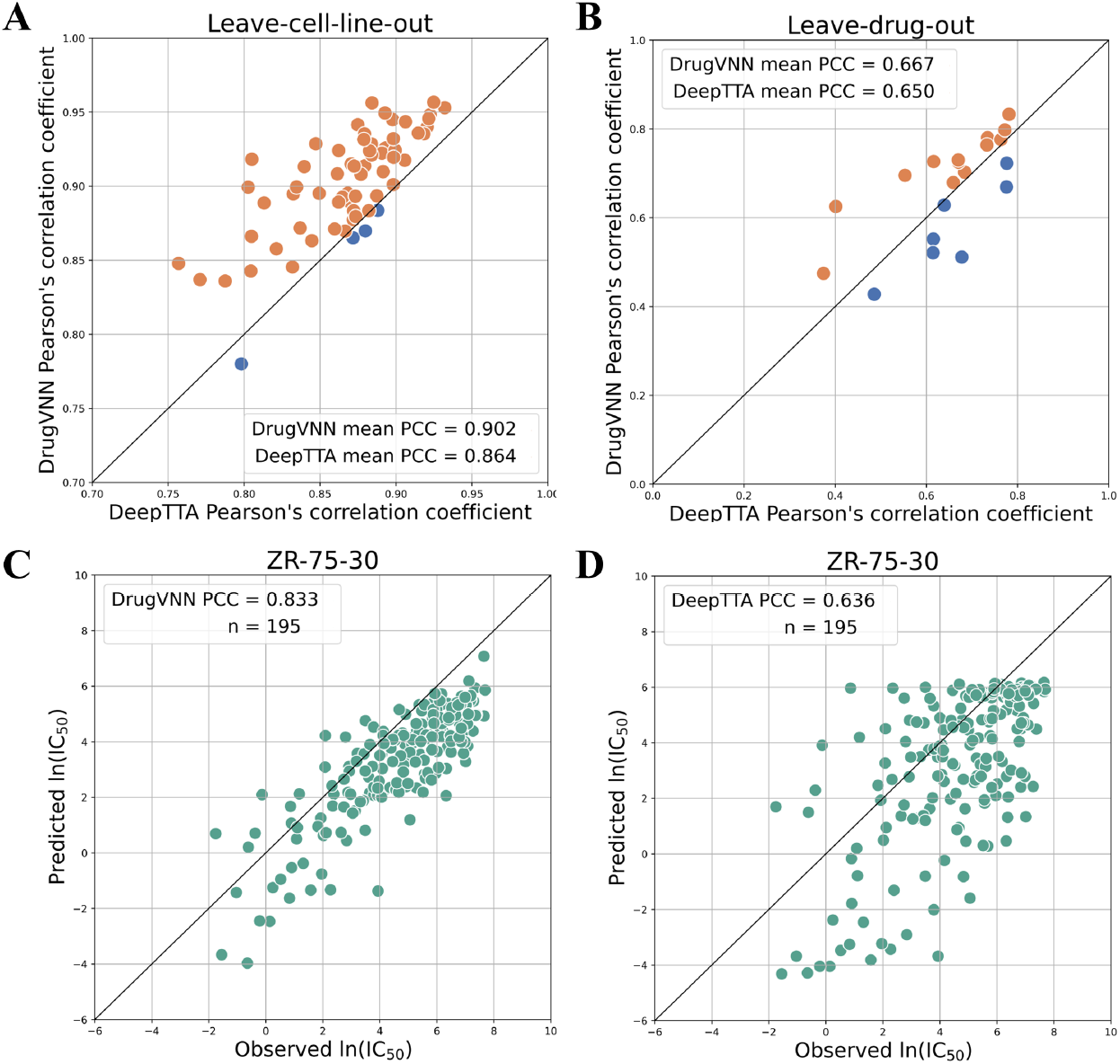
Comparison of PCC between DrugVNN and DeepTTA in the leave-cell-out and leave-drug-out scenario (A and B), respectively. The dot is orange if DrugVNN’s PCC is higher than DeepTTA, otherwise is blue. C and D illustrate a prediction case of DrugVNN and DeepTTA in cell line ZR-75-30.

**Fig. 3.**
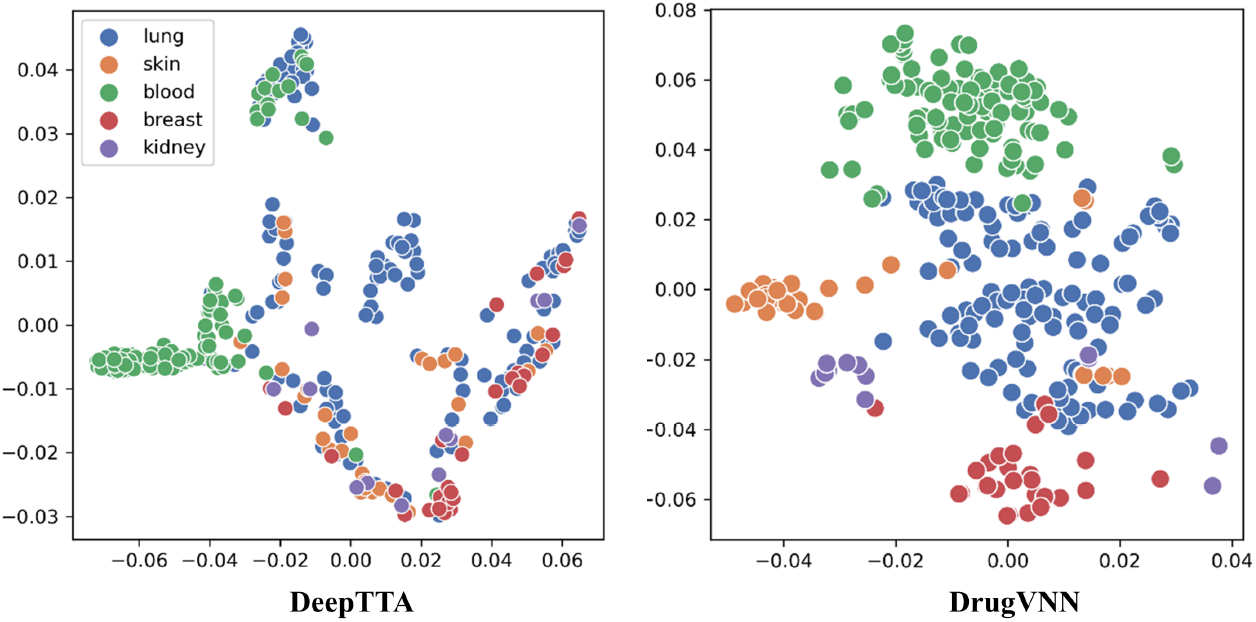
The t-SEN projection of cell line representation learned from DeepTTA and DrugVNN colored by corresponding tissue.

### 3.3. Predicting binary drug sensitivity status

To explore the predictive power in classifying drug sensitivity of DrugVNN, we binarized IC_50_ according to the threshold of each drug provided by [34]. After filtering drug samples without a binary threshold, we collected a dataset with 30,505 positive instances in which drugs are sensitive to the corresponding cell lines and 2,800 negative instances where drugs are resistant. We compared DrugVNN to three other neural network models by rediscovering the drug response status. Despite the unbalanced dataset, DrugVNN outperforms all other methods by achieving a higher AUC and AUPR score of 0.907 and 0.543 (Fig. 4), reaffirming the advanced predictive performance of DrugVNN.

**Fig. 4.**
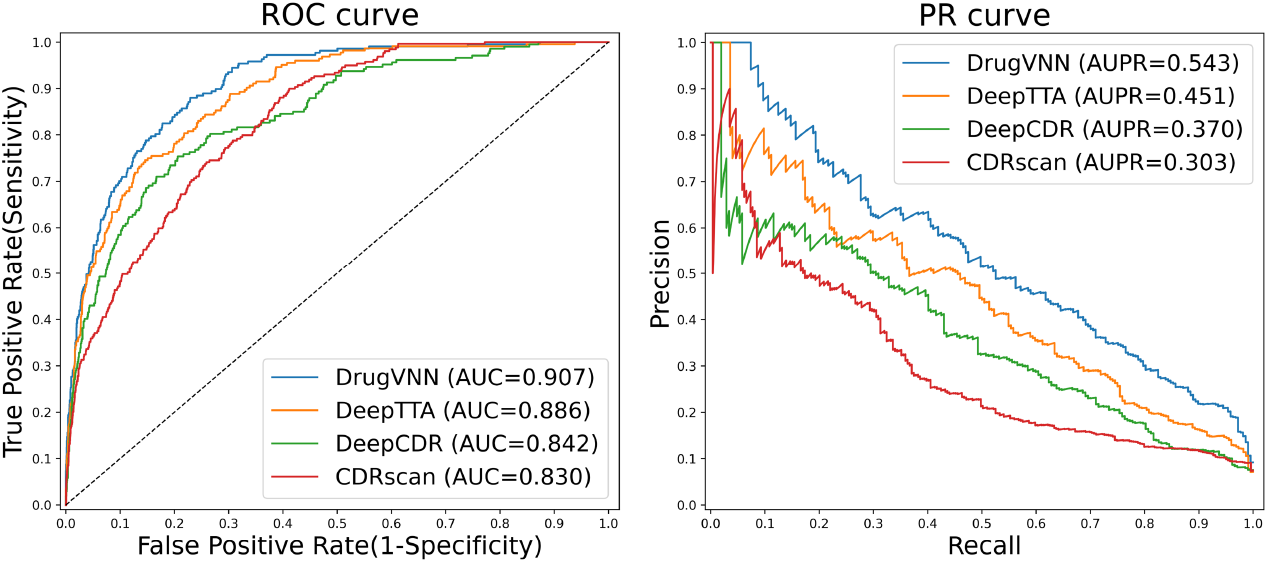
Comparision of the ROC and PR curve between DrugVNN and baselines.

### 3.4. Model components analysis

To study the contributions of each module of DrugVNN, we conducted ablation studies from two perspectives: drug features and cell line features encoder. Firstly, we replaced the CMPNN with 3 commonly used GNN models: GCN, SAGE and GIN. As shown in the upper half of Table 2, DrugVNN surpassed all three GNN variants. The advancement should be attributed to the strengthened communications between nodes and edges in DrugVNN, while other models only focused on the node-to-node message passing mechanism.

**Table 2.** Ablation studies of DrugVNN.

|  |  | RMSE | MAE | R<sup>2</sup> |
| --- | --- | --- | --- | --- |
| Drug | SAGE | 0.952 | 0.700 | 0.879 |
|  | GCN | 0.949 | 0.699 | 0.879 |
|  | GIN | 0.945 | 0.697 | 0.881 |
| Cell line | FCN | 0.981 | 0.739 | 0.872 |
|  | Random VNN | 1.055 | 0.781 | 0.853 |
|  | w/o drug-aware attention | 0.957 | 0.712 | 0.875 |
|  | DrugVNN | **0.935** | **0.685** | **0.885** |

To assess the effects of visible neural network, we replaced the VNN with the matched fully connected network which has the same number of neurons in each hidden layer and the same depth as DrugVNN. We also generated a random VNN by shuffling the pathway-pathway relationship as a control. As shown in the lower half of Table 2, when adopting the fully connected network, the model showed poor performance with a 0.046 increase in RMSE and a 0.013 drop in R^2^, and the performance would further degrade when adopting the random VNN. These results demonstrated that the non-random gene pathways profile information could facilitate the learning of cell line representation for precise prediction. Moreover, we also disabled the drug-aware gene attention gates, which caused a 0.022 increase in RMSE, demonstrating that the attention mechanism can enhance the learning ability of VNN.

### 3.5. Case Studies of DrugVNN

#### 3.5.1. Missing drug-cell line responses discovery

Missing values prediction has been widely used in drug response prediction studies to identify whether the model is capable of inductive prediction. We applied our model to predict the missing IC_50_ values and sorted the average IC_50_ value for each drug. Fig. 5 illustrates the distributions of the predicted values. Daporinad, Docetaxel and Bortezomib are the top-3 sensitive drugs, which is consistent with the prediction of previous DRP methods[23, 24, 35]. The most sensitive drug, Daporinad, is one of the highly specific inhibitors of nicotinamide phosphoribosyl transferase and known to have its unique mechanism of action that induces the tumor cell apoptosis[36]. Among the prediction results of Daporinad, (Daporinad, MOLM-13) has the lowest value of −8.82. MOLM-13 cell line is established from the peripheral blood of a patient at relapse of acute monocytic leukemia[37]. We also found that 9 of the top-10 sensitive cell lines of Daporinad are both associated with leukemia (Supplementary Table S4). These findings are consistent with the report that most hematologic cancer cells are sensitive to low concentrations of Daporinad as measured in cytotoxicity and clonogenic assays[38].

**Fig. 5.**
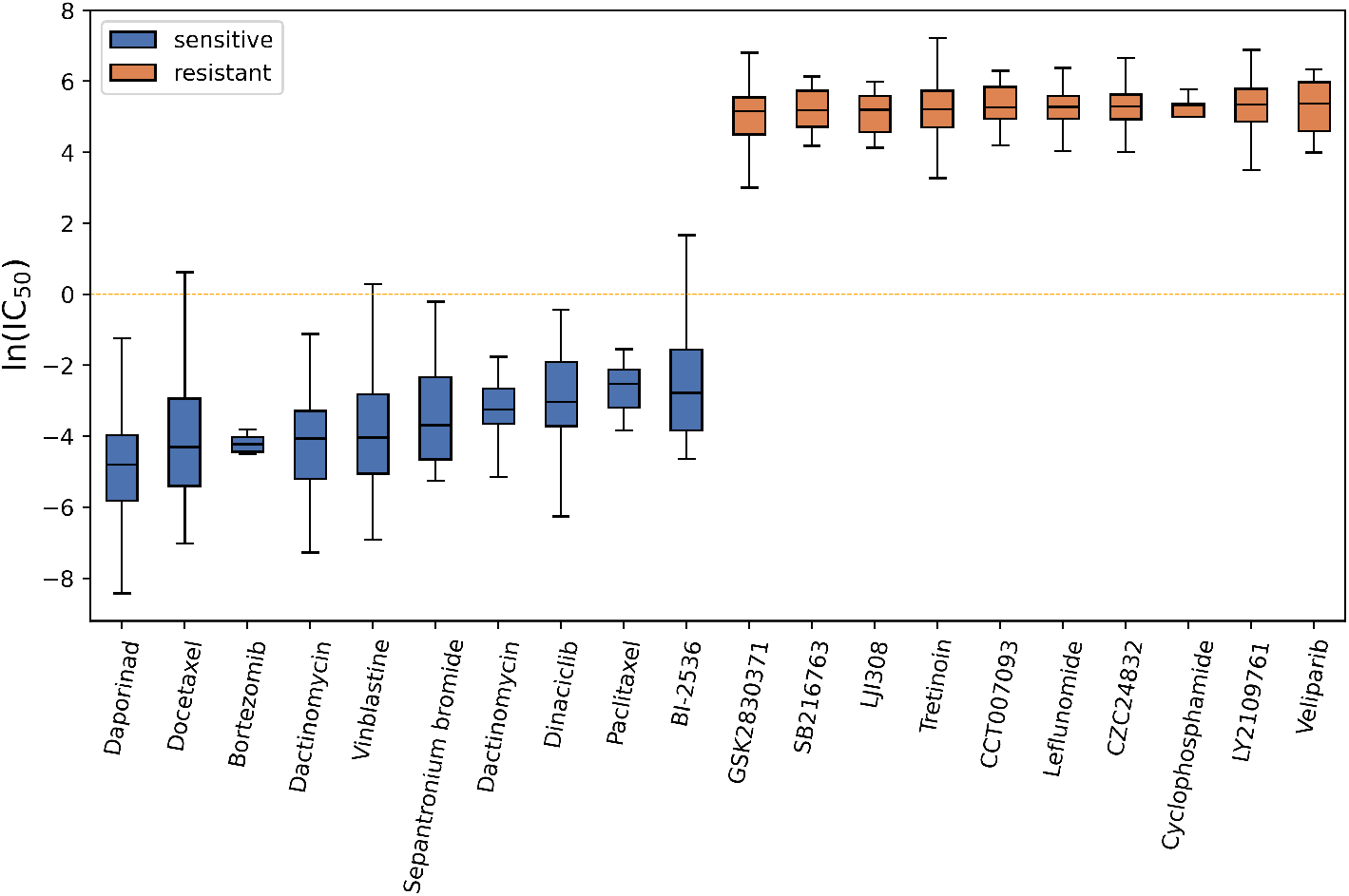
Top-10 sensitivity and resistance drugs of the missing drug responses predicted by DrugVNN.

#### 3.5.2. Gene-level interpretation through drug-aware gene attention gates

To determine whether DrugVNN could capture meaningful gene information, we performed gene-level analysis through the drug-aware gene attention gates. Specifically, for each drug-cell line pair in the test set, the genes with normalized attention scores above 0.99 are selected as the potential target genes for the input drug. The ground-truth drug-target interactions are downloaded from STITCH database[39]. Among the 1,626 predicted drug-target pairs, 220 (13.5%) were reported in STITCH, which is significantly higher than 18.6 ± 3.1 drug-target pairs from the control when replacing the learned attention matrix with a randomly generated matrix (*P* = 1.1 × 10^−4^ by two-sided *t* − *test*). Table 3 listed the top-5 important genes of four drug-cell line pairs found by DrugVNN, with target genes obtained from STITCH labeled as bold. These results suggested that our method could successfully identify and pay more attention to the putative key genes, and thus predict the drug response more precisely.

**Table 3.**
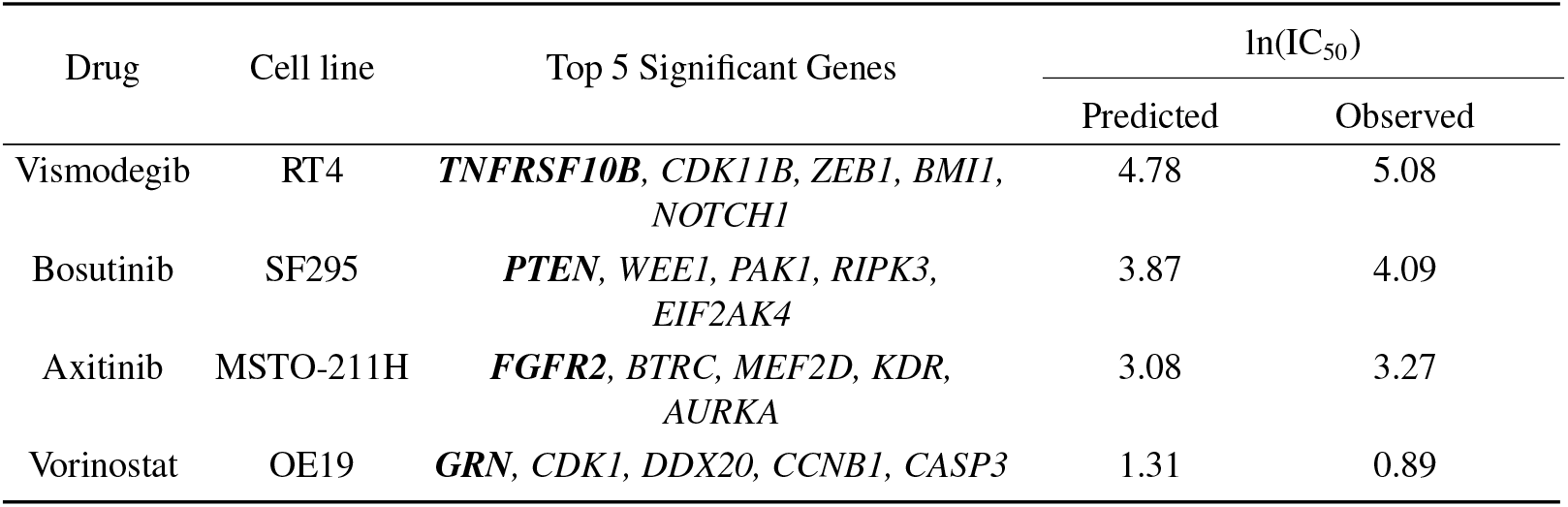
Top-5 important genes detected by DrugVNN with respect to drug-cell line pairs. Genes in bold are targets of drugs.

#### 3.5.3. Pathway-level interpretation through knowledge-guided visible network

To understand the biological process inside cells that contributed to the predictive performance, we visualized the high-attribution sub-network structure of DrugVNN. Specifically, we selected the drug-cell line pair (Pictilisib, A253) as a case to study the path of impact from the input to the outcome. As shown in Fig. 6, the path with the highest SHAP values begin with a gene, PIK3CA, connected to a hierarchy of pathways including constitutive signaling by aberrant PI3K in cancer, PI3K/AKT signaling in cancer, and diseases of signal transduction by the growth factor receptors. Pictilisib was reported to bind to the adenosine triphosphate-binding pocket of PI3K to prevent the formation of phosphatidylinositol (3,4,5)-trisphosphate, a key component of the PI3K pathways, and thus finally selectively inhibits class I PI3K[40]. The PI3K signaling pathway is a key regulator of tumor proliferation and mutations of PIK3CA have been identified in many kinds of cancers[41, 42]. Analysis of 151 head and neck squamous cell carcinoma (HNSCC) cell lines reveals PI3K pathway was the most frequently mutated oncogenic pathway (30.5%) of HNSCC compared to other known cancer genes[43]. A253, from the submaxillary salivary gland, is one kind of HNSCC. The activation of PI3K alterations happens in the majority of salivary duct carcinoma (SDC) patients, and PI3K targeted therapy proved to be a potential combination in patients with SDC[44]. This complete interpretation of drug-pathways-genes demonstrates the ability of DrugVNN to identify the genes and pathways related to the drug’s target, thus promoting practical clinical cancer drug discovery.

**Fig. 6.**
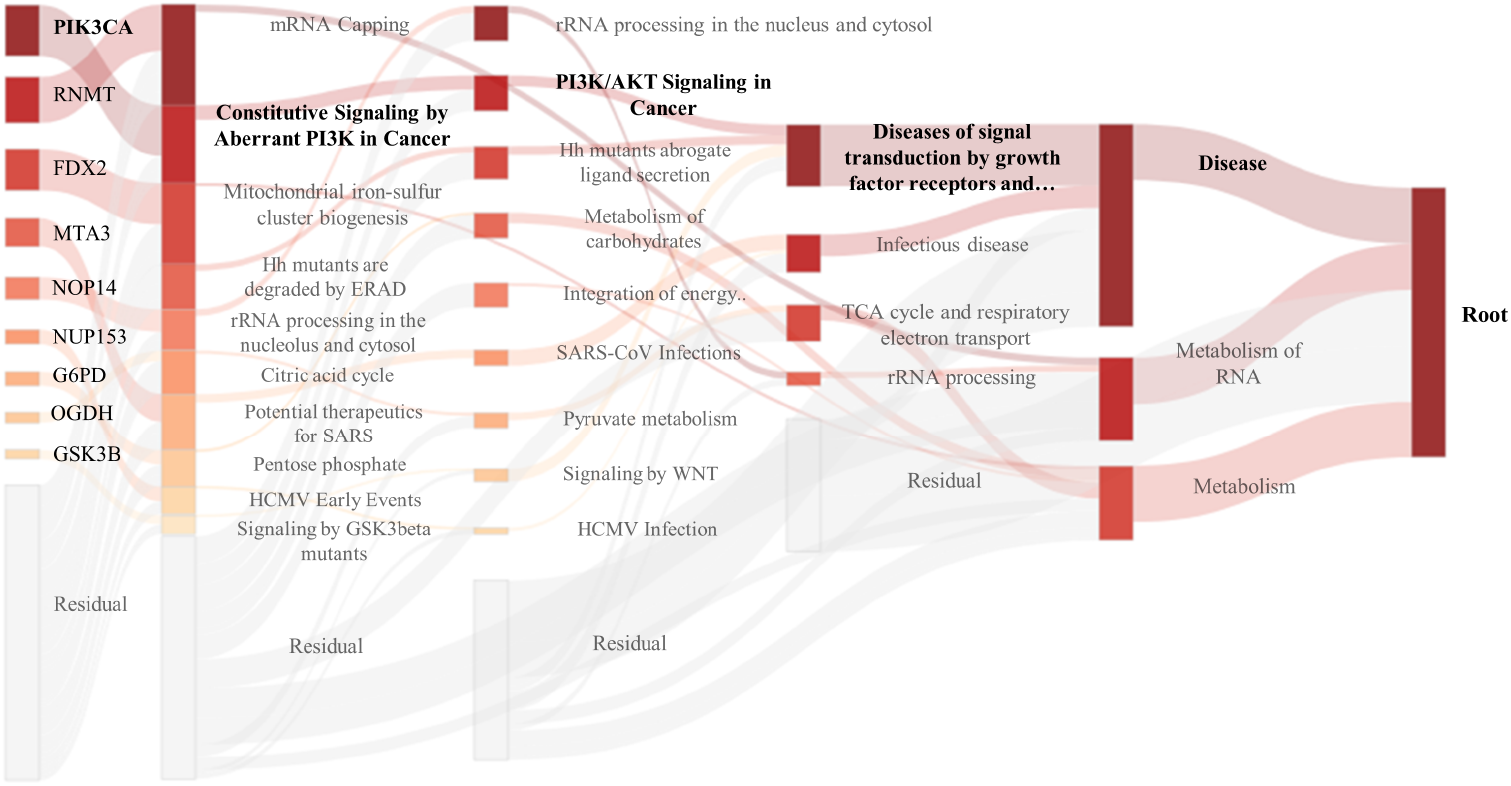
Visualization of inner layers of DrugVNN shows the estimated relative importance of different nodes in each layer. Nodes on the far left represent input genes and nodes in the middle layers represent pathways. Nodes with darker colors are more important, while transparent nodes represent the undisplayed nodes in each layer.

## 4. Conclusion and future directions

Different cancer patients may respond differently to cancer treatment due to the complexity and heterogeneity of cancer, leading to a growing demand to develop an efficient computational method for identifying drug responses in cancer cell lines. In this study, we proposed a novel drug-response prediction model, DrugVNN, providing predictive performance as well as interpretability. We incorporated node-edge communicative message passing network and hierarchical pathway information-guided visible network to extract drug and cell lien features. In addition, we designed a novel drug-aware gene attention gate mechanism, which selected important genes for the given drug and optimized cell line representation learning. The performance comparison experiments indicated that DrugVNN has demonstrated superior predictive power when compared to the state-of-the-art DRP models. Furthermore, we implemented gene- and pathway-level analysis via DrugVNN and delineated the mechanism governing the drug response under specific drug-cell line pairs.

However, there is still room for further improvements on DrugVNN. First of all, pre-training strategy has been widely adopted in molecule representation learning[45]. The performance of DRP could be improved if an appropriate task-related pre-training method can be applied. Secondly, we only employed the attention gate mechanism on the very first layer of the VNN considering training costs. In fact, drug information can be added to each layer to better enhance the representation learning of the cell line. In the future, we would integrate more prior knowledge, such as cell painting[46] and knowledge graphs[47], to further improve our model and expand the framework to other applications, such as drug combination prediction, survival analysis[48], etc.

## References

[1] R. L. Siegel, K. D. Miller, N. S. Wagle, A. Jemal, Cancer statistics, 2023, Ca Cancer J Clin 73 (1) (2023) 17–48.

[2] C. H. Wong, K. W. Siah, A. W. Lo, Estimating clinical trial success rates and related parameters in oncology, Available at SSRN 3355022 (2019).

[3] W. Lu, Q. Wu, J. Zhang, J. Rao, C. Li, S. Zheng, Tankbind: Trigonometry-aware neural networks for drug-protein binding structure prediction, Advances in neural information processing systems 35 (2022) 7236–7249.

[4] Y.-C. Chiu, H.-I. H. Chen, A. Gorthi, M. Mostavi, S. Zheng, Y. Huang, Y. Chen, Deep learning of pharmacogenomics resources: moving towards precision oncology, Briefings in bioinformatics 21 (6) (2020) 2066–2083.

[5] W. Yang, J. Soares, P. Greninger, E. J. Edelman, H. Lightfoot, S. Forbes, N. Bindal, D. Beare, J. A. Smith, I. R. Thompson, et al., Genomics of drug sensitivity in cancer (gdsc): a resource for therapeutic biomarker discovery in cancer cells, Nucleic acids research 41 (D1) (2012) D955–D961.

[6] J. Barretina, G. Caponigro, N. Stransky, K. Venkatesan, A. A. Margolin, S. Kim, C. J. Wilson, J. Lehar, G. V. Kryukov, D. Sonkin, et al., The cancer cell line encyfclopedia enables predictive modelling of anticancer drug sensitivity, Nature 483 (7391) (2012) 603–607.

[7] D. Baptista, P. G. Ferreira, M. Rocha, Deep learning for drug response prediction in cancer, Briefings in bioinformatics 22 (1) (2021) 360–379.

[8] M. P. Menden, F. Iorio, M. Garnett, U. McDermott, C. H. Benes, P. J. Ballester, J. Saez-Rodriguez, Machine learning prediction of cancer cell sensitivity to drugs based on genomic and chemical properties, PLoS one 8 (4) (2013) e61318.

[9] Y. Wang, J. Fang, S. Chen, Inferences of drug responses in cancer cells from cancer genomic features and compound chemical and therapeutic properties, Scientific reports 6 (1) (2016) 32679.

[10] L. Wang, X. Li, L. Zhang, Q. Gao, Improved anticancer drug response prediction in cell lines using matrix factorization with similarity regularization, BMC cancer 17 (1) (2017) 1–12.

[11] Y. Chang, H. Park, H.-J. Yang, S. Lee, K.-Y. Lee, T. S. Kim, J. Jung, J.-M. Shin, Cancer drug response profile scan (cdrscan): a deep learning model that predicts drug effectiveness from cancer genomic signature, Scientific reports 8 (1) (2018) 8857.

[12] H. Sharifi-Noghabi, O. Zolotareva, C. C. Collins, M. Ester, Moli: multi-omics late integration with deep neural networks for drug response prediction, Bioinformatics 35 (14) (2019) i501–i509.

[13] P. Liu, H. Li, S. Li, K.-S. Leung, Improving prediction of phenotypic drug response on cancer cell lines using deep convolutional network, BMC bioinformatics 20 (1) (2019) 1–14.

[14] J. Rao, S. Zheng, Y. Lu, Y. Yang, Quantitative evaluation of explainable graph neural networks for molecular property prediction, Patterns 3 (12) (2022).

[15] G. Eraslan, Z. Avsec, J. Gagneur, F. J. Theis, Deep learning: new computational modelling techniques for genomics, Nature Reviews Genetics 20 (7) (2019) 389–403.

[16] K. Y. Michael, J. Ma, J. Fisher, J. F. Kreisberg, B. J. Raphael, T. Ideker, Visible machine learning for biomedicine, Cell 173 (7) (2018) 1562–1565.

[17] B. M. Kuenzi, J. Park, S. H. Fong, K. S. Sanchez, J. Lee, J. F. Kreisberg, J. Ma, T. Ideker, Predicting drug response and synergy using a deep learning model of human cancer cells, Cancer cell 38 (5) (2020) 672–684.

[18] L. Deng, Y. Cai, W. Zhang, W. Yang, B. Gao, H. Liu, Pathway-guided deep neural network toward interpretable and predictive modeling of drug sensitivity, Journal of Chemical Information and Modeling 60 (10) (2020) 4497–4505.

[19] G. O. Consortium, Expansion of the gene ontology knowledgebase and resources, Nucleic acids research 45 (D1) (2017) D331–D338.

[20] M. Kramer, J. Dutkowski, M. Yu, V. Bafna, T. Ideker, Inferring gene ontologies from pairwise similarity data, Bioinformatics 30 (12) (2014) i34–i42.

[21] C. W. Coley, R. Barzilay, W. H. Green, T. S. Jaakkola, K. F. Jensen, Convolutional embedding of attributed molecular graphs for physical property prediction, Journal of chemical information and modeling 57 (8) (2017) 1757–1772.

[22] T. Nguyen, G. T. Nguyen, T. Nguyen, D.-H. Le, Graph convolutional networks for drug response prediction, IEEE/ACM transactions on computational biology and bioinformatics 19 (1) (2021) 146–154.

[23] Q. Liu, Z. Hu, R. Jiang, M. Zhou, Deepcdr: a hybrid graph convolutional network for predicting cancer drug response, Bioinformatics 36 (Supplement 2) (2020) i911–i918.

[24] L. Jiang, C. Jiang, X. Yu, R. Fu, S. Jin, X. Liu, Deeptta: a transformer-based model for predicting cancer drug response, Briefings in bioinformatics 23 (3) (2022) bbac100.

[25] Y. Song, S. Zheng, Z. Niu, Z.-H. Fu, Y. Lu, Y. Yang, Communicative representation learning on attributed molecular graphs., in: IJCAI, Vol. 2020, 2020, pp. 2831–2838.

[26] J. Rao, S. Zheng, S. Mai, Y. Yang, Communicative subgraph representation learning for multi-relational inductive drug-gene interaction prediction, arXiv preprint arXiv:2205.05957 (2022).

[27] A. Vaswani, N. Shazeer, N. Parmar, J. Uszkoreit, L. Jones, A. N. Gomez, Ł. Kaiser, I. Polosukhin, Attention is all you need, Advances in neural information processing systems 30 (2017).

[28] Y. Wang, J. Xiao, T. O. Suzek, J. Zhang, J. Wang, S. H. Bryant, Pubchem: a public information system for analyzing bioactivities of small molecules, Nucleic acids research 37 (suppl 2) (2009) W623–W633.

[29] A. Fabregat, S. Jupe, L. Matthews, K. Sidiropoulos, M. Gillespie, P. Garapati, R. Haw, B. Jassal, F. Korninger, B. May, et al., The reactome pathway knowledgebase, Nucleic acids research 46 (D1) (2018) D649–D655.

[30] J. Gilmer, S. S. Schoenholz, P. F. Riley, O. Vinyals, G. E. Dahl, Message passing neural networks, Machine learning meets quantum physics (2020) 199–214.

[31] S. M. Lundberg, S.-I. Lee, A unified approach to interpreting model predictions, Advances in neural information processing systems 30 (2017).

[32] M. Sun, S. Zhao, C. Gilvary, O. Elemento, J. Zhou, F. Wang, Graph convolutional networks for computational drug development and discovery, Briefings in bioinformatics 21 (3) (2020) 919–935.

[33] L. Van der Maaten, G. Hinton, Visualizing data using t-sne., Journal of machine learning research 9 (11) (2008).

[34] F. Iorio, T. A. Knijnenburg, D. J. Vis, G. R. Bignell, M. P. Menden, M. Schubert, N. Aben, E. Goncalves, S. Barthorpe, H. Lightfoot, et al., A landscape of pharmacogenomic interactions in cancer, Cell 166 (3) (2016) 740–754.

[35] Y. Zhu, Z. Ouyang, W. Chen, R. Feng, D. Z. Chen, J. Cao, J. Wu, Tgsa: protein–protein association-based twin graph neural networks for drug response prediction with similarity augmentation, Bioinformatics 38 (2) (2022) 461–468.

[36] M. Park, B. I. Lee, J. Choi, Y. Park, S.-J. Park, J.-H. Lim, J. Lee, Y. G. Shin, Quantitative analysis of daporinad (fk866) and its in vitro and in vivo metabolite identification using liquid chromatography-quadrupole-time-of-flight mass spectrometry, Molecules 27 (6) (2022) 2011.

[37] Y. Matsuo, R. MacLeod, C. Uphoff, H. Drexler, C. Nishizaki, Y. Katayama, G. Kimura, N. Fujii, E. Omoto, M. Harada, et al., Two acute monocytic leukemia (aml-m5a) cell lines (molm-13 and molm-14) with interclonal phenotypic heterogeneity showing mll-af9 fusion resulting from an occult chromosome insertion, ins (11; 9)(q23; p22p23), Leukemia 11 (9) (1997) 1469–1477.

[38] A. Nahimana, A. Attinger, D. Aubry, P. Greaney, C. Ireson, A. V. Thougaard, J. Tjørnelund, K. M. Dawson, M. Dupuis, M. A. Duchosal, The nad biosynthesis inhibitor apo866 has potent antitumor activity against hematologic malignancies, Blood, The Journal of the American Society of Hematology 113 (14) (2009) 3276–3286.

[39] D. Szklarczyk, A. Santos, C. Von Mering, L. J. Jensen, P. Bork, M. Kuhn, Stitch 5: augmenting protein–chemical interaction networks with tissue and affinity data, Nucleic acids research 44 (D1) (2016) D380–D384.

[40] A. J. Folkes, K. Ahmadi, W. K. Alderton, S. Alix, S. J. Baker, G. Box, I. S. Chuckowree, P. A. Clarke, P. Depledge, S. A. Eccles, et al., The identification of 2-(1 h-indazol-4-yl)-6-(4-methanesulfonyl-piperazin-1-ylmethyl)-4-morpholin-4-yl-thieno [3, 2-d] pyrimidine (gdc-0941) as a potent, selective, orally bioavailable inhibitor of class i pi3 kinase for the treatment of cancer, Journal of medicinal chemistry 51 (18) (2008) 5522–5532.

[41] C. Porta, C. Paglino, A. Mosca, Targeting pi3k/akt/mtor signaling in cancer, Frontiers in oncology 4 (2014) 64.

[42] W.-C. Huang, M.-C. Hung, Induction of akt activity by chemotherapy confers acquired resistance, Journal of the Formosan Medical Association 108 (3) (2009) 180–194.

[43] V. W. Lui, M. L. Hedberg, H. Li, B. S. Vangara, K. Pendleton, Y. Zeng, Y. Lu, Q. Zhang, Y. Du, B. R. Gilbert, et al., Frequent mutation of the pi3k pathway in head and neck cancer defines predictive biomarkers, Cancer discovery 3 (7) (2013) 761–769.

[44] P. Saintigny, Y. Mitani, K. B. Pytynia, R. Ferrarotto, D. B. Roberts, R. S. Weber, M. S. Kies, S. N. Maity, S.-H. Lin, A. K. El-Naggar, Frequent pten loss and differential her2/pi3k signaling pathway alterations in salivary duct carcinoma: Implications for targeted therapy, Cancer 124 (18) (2018) 3693–3705.

[45] B. Fabian, T. Edlich, H. Gaspar, M. Segler, J. Meyers, M. Fiscato, M. Ahmed, Molecular representation learning with language models and domain-relevant auxiliary tasks, arXiv preprint arXiv:2011.13230 (2020).

[46] M.-A. Bray, S. Singh, H. Han, C. T. Davis, B. Borgeson, C. Hartland, M. Kost-Alimova, S. M. Gustafsdottir, C. C. Gibson, A. E. Carpenter, Cell painting, a high-content image-based assay for morphological profiling using multiplexed fluorescent dyes, Nature protocols 11 (9) (2016) 1757–1774.

[47] S. Zheng, J. Rao, Y. Song, J. Zhang, X. Xiao, E. F. Fang, Y. Yang, Z. Niu, Pharmkg: a dedicated knowledge graph benchmark for bomedical data mining, Briefings in bioinformatics 22 (4) (2021) bbaa344.

[48] Y. Wang, Z. Zhang, H. Chai, Y. Yang, Multi-omics cancer prognosis analysis based on graph convolution network, in: 2021 IEEE International Conference on Bioinformatics and Biomedicine (BIBM), IEEE, 2021, pp. 1564–1568.

